# Somalier: rapid relatedness estimation for cancer and germline studies using efficient genome sketches

**DOI:** 10.1101/839944

**Authors:** Brent S Pedersen, Preeti J Bhetariya, Joe Brown, Gabor Marth, Randy L Jensen, Mary P Bronner, Hunter R Underhill, Aaron R Quinlan

## Abstract

When interpreting sequencing data from multiple spatial or longitudinal biopsies, detecting sample mix-ups is essential yet more difficult than in studies of germline variation. In most genomic studies of tumors, genetic variation is frequently detected through pairwise comparisons of the tumor and a matched normal tissue from the sample donor, and in many cases, only somatic variants are reported. The disjoint genotype information that results hinders the use of existing tools that detect sample swaps solely based on genotypes of germline variants. To address this problem, we have developed *somalier*, which can operate directly on the alignments, so as not to require jointly-called germline variants. Instead, *somalier* extracts a small *sketch* of informative genetic variation for each sample. Sketches from hundreds of biopsies and normal tissues can then be compared in under a second. This speed also makes it useful for checking relatedness in large cohorts of germline samples. Somalier produces both text output and an interactive visual report that facilitates the detection and correction of sample swaps using multiple relatedness metrics. We introduce the tool and demonstrate its utility on a cohort of five glioma samples each with a normal, tumor, and cell-free DNA sample. Applying *somalier* to high-coverage sequence data from the 1000 Genomes Project also identifies several related samples. *Somalier* can be applied to diverse sequencing data types and genome builds, and is freely available for academic use at github.com/brentp/somalier.

## Introduction

DNA sequencing data from matched tumor-normal pairs are critical for the detection of somatic variation in cancer studies. However, a sample swap leads to a dramatic increase in the apparent number of somatic variants, confounds the genetic analysis of the tumor, and the probability of such a mix-up increases with the size of the study cohort. The correction for sample mix-ups, possibly a swap with another sample in the same study, requires a thorough evaluation of relatedness among the entire set of samples. This is not possible directly on the somatic mutation predictions because somatic variants are typically called as tumor-normal pairs, and often, only somatic (not germline) variants are reported^1^. Therefore, resolution of the sample swap would require the researcher to jointly call germline variants with a tool like GATK^2^ and then use methods such as peddy^3^ or KING^4^ to calculate relatedness across the entire set of samples. Joint variant-calling is time and resource intensive, especially when all that is needed to resolve the sample swap is an accurate calculation of relatedness among the samples. After experiencing this inconvenience in our own research, we developed *somalier* to quickly and accurately compute relatedness by extracting “sketches” of variant information directly from alignments (BAM or CRAM) or from variant-call format (VCF)^5^ files including genomic VCFs (GVCF). *Somalier* extracts a sketch for each sample and the sketches are then compared to evaluate all possible pairwise relationships among the samples. This setup allows users to add new sketches as needed and then efficiently compare to a set of background samples. The text and visual output facilitates the detection and correction of sample swaps, even in cases where there is severe loss-of-heterozygosity. It can be used on any organism across diverse sequencing data types, and, given a set of carefully selected sites, across genome builds.

## Methods

### Selecting and extracting informative variant sites

We have previously shown that using as few as 5,000 carefully chosen polymorphic loci is sufficient for relatedness estimation, is more accurate than using all available variants^3^. A similar site-selection strategy is also used in Conpair to estimate contamination^6^. Using the fewest and best sites enables fast, yet very accurate relatedness estimation. In *somalier*, we utilize the observation that the optimal sites for detecting relatedness are high-quality, unlinked sites with an allele frequency of around 0.5. A balanced allele frequency maximizes the probability that any 2 unrelated samples will differ. We distribute a set of sites to be queried by *somalier*. The sites are high-frequency single-nucleotide variants selected from gnomAD^7^ exomes that exclude segmental duplication and low-complexity regions^8^. Variants with nearby insertions or deletions are excluded. In addition, we have excluded sites that are cytosines in the reference so that the tool can be used on bisulfite seq data, for example, to check the correspondence between bisulfite sequencing and RNA-Seq data. The *somalier* repository includes the code to create a set of sites for different organisms given a population VCF and a set of optional exclude regions. We distribute a set of matched sites for both the GRCh37 and GRCh38 builds of the human reference genome; that is, a user can extract sites from a sample aligned to GRCh37 using our GRCh37 sites file and compare that sketch to a sketch created from a sample aligned to GRCh38 by extracting the sites in our GRCh38 file. This is convenient as labs move from GRCh37 to GRCh38 and future genome builds. The sites files include informative variants on the X and Y chromosomes so that *somalier* can also estimate a sample’s sex from the genotypes. However, only autosomal sites are used to estimate relatedness. In total, somalier inspects 17,766 total sites (these are distributed with the somalier software and available as Supplementary Files 1 and 2), all of which are chosen to be in coding sequence so that they are applicable to genome, exome, and RNA-seq datasets.

In order to quickly extract data from polymorphic sites into a genome sketch, *somalier* uses the BAM or CRAM index to query each file at each of the sites described above. Alignments with a mapping quality of at least 1 that are not duplicates, supplementary, or failing quality control (according to the SAM flag) are used. The CIGAR string of each passing alignment is evaluated at the requested position and the base in the alignment at that position is checked against the given reference and alternate for the query variant. This is faster than a traditional sequence alignment “pileup” as it looks at each read only once and interrogates only the exact position in question. If a VCF (or BCF or GVCF) is provided instead of an alignment file, *somalier* will extract the depth information for each sample for requested sites that are present in the VCF. The sketches extracted from a VCF are indistinguishable from those extracted from alignment files. In order to support single-sample VCFs, which do not contain homozygous reference calls, a user can indicate that missing variants should be assumed to be homozygous reference. This also facilitates comparing multiple tumor-normal VCFs where many sites will not be shared (however, in those cases, it’s preferable extract the sketch from the alignment files rather than from the VCF).

*Somalier* tallies reference and alternate counts for each site. Once all sites are collected, it writes a binary file containing the sample name and the allele counts collected at each of the inspected sites. For the set of sites distributed from the somalier repository, a sketch files requires ~200KB of space on disk or in memory. This sketch format and the speed of parsing and comparing sketch files are key strengths of *somalier*. For example, since *somalier* can complete a full analysis of 2504 sketches from the 1000 Genomes high-coverage whole-genome samples (Michael Zody, personal communication) in under 20 seconds, users can keep a pool of sample sketches to test against and check incoming samples against all previously sketched samples.

### Comparing Sketches

Thousands of sample sketches can be read into memory per second and compared. In order to calculate relatedness, *somalier* converts the ref and alt counts stored for each sample at each site into a genotype. The genotype is determined to be unknown if the depth is less than the user-specified value (default of 7), homozygous reference if the allele balance (i.e., alt-count / [ref-count + alt-count]) is less than 0.02, heterozygous if the allele balance is between 0.2 and 0.8, homozygous alternate if the allele balance is above 0.98 and unknown otherwise (**Figure 1A**). A flag can amend these rules such that missing sites (with depth of 0) are treated as homozygous reference, rather than unknown.

**Figure 1.**
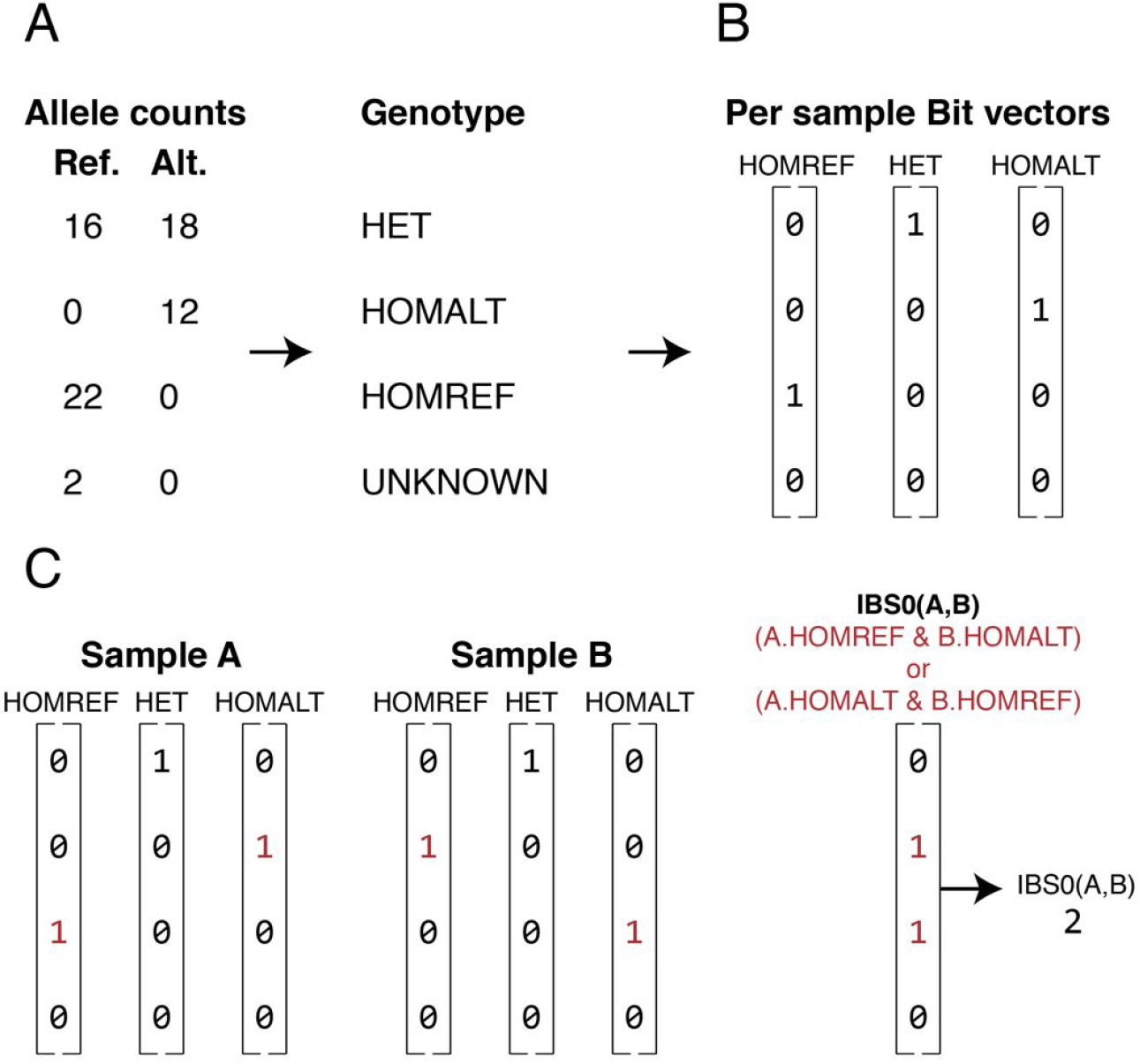
Comparing genotype sketches to compute relatedness measures for pairs of samples. **A.** Observed counts for the reference (Ref.) and alternate (Alt.) allele at each of the tested 17,766 loci are converted into genotypes (see main text for details) to create a “sketch” for each sample. **B.** The genotypes for each sample are then converted into three bit vectors: one for homozygous reference (HOMREF) genotypes, one for heterozygous (HET) genotypes, and one for homozygous alternate (HOMALT) genotypes. The length of each vector is the total number of autosomal variants in the sketch (i.e., 17,384) divided by 64, and the value for each bit is set to 1 if the sample has the particular genotype at the given variant site. For example, four variant sites are shown in panel B and the hypothetical individual has a homozygous alternate genotype for the second variant (the corresponding bit is set to 1), but is not homozygous for the alternate allele at the other three variant sites (the corresponding bits are set to 0). **C.** The bit vectors for a pair of samples can be easily compared to calculate relatedness measures (e.g., IBS0) through efficient, bitwise operations on the bit arrays for the relevant genotypes.

While simple, this heuristic genotyping works well in practice and is extremely fast, because *somalier* looks only at single-nucleotide variants in non-repeat regions of the genome. As the sample is processed, *somalier* also collects information on depth, mean allele-balance, number of reference, heterozygous, and homozygous alternate calls for each sample, along with similar stats for the X and Y chromosomes. These data are used to calculate per-sample quality control metrics. In order to measure relatedness, the data collected for each sample is converted into a data structure consisting of *hom_ref, het*, and *hom_alt* bit vectors (**Figure 1B**). The bit vectors consist of 64 bit integers, enabling *somalier* to store 64 variants per integer. There are 17,384 autosomal sites in the default sites file used by *somalier*, consuming only 6519 bytes per sample (17384 / 64 bits * 3 bit-vectors/sample * 8 bits/byte). With this data layout, *somalier* can represent all 2,504 samples from the 1000 Genomes Project in under 17 megabytes of memory. This simple data structure also facilitates rapid pairwise comparisons (**Figure 1C**); for example, we can compute IBS0 (that is, “identity by-state 0” or sites where zero alleles are shared between two samples *A* and *B*) with the following logic which evaluates 64 sites in parallel:

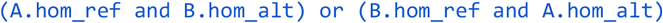

We repeat this for each of the 272 (17384 autosomal sites / 64 sites per entry) entries in the array to assess all of the genome-wide sites for each pair of samples. In fact, we don’t need the sites, just the count of sites that are IBS0. Therefore, we use the *popcount* (i.e. the count of bits that are set to TRUE) hardware instruction to count the total number of bits where the expression is non-zero in order to get the total count of IBS0 sites between the 2 samples. In addition to IBS0, we calculate counts of IBS2 where both samples have the same genotype, shared heterozygotes (both are heterozygotes), shared homozygous alternates, and heterozygous sites for each sample. All of the operations are extremely fast as it does not require code branching via, for example, conditional logic; instead the calculations are all conducted with bitwise operations.

Once those metrics are calculated, the relatedness between sample *i* and sample *j* is calculated as:

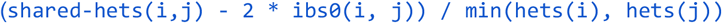

where *hets* is the count of heterozygote calls per sample out of the assayed sites. In addition, the *homozygous concordance* rate is reported as:

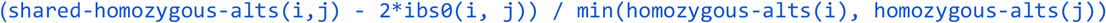

This measure is similar to the one described in HYSYS9 except that the HYSYS measure is simply:

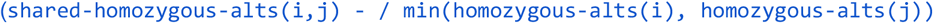

Our formulation has the benefit that it matches the scale and interpretation of the relatedness estimate; unrelated individuals will have a concordance of around 0, whereas in HYSYS they will have a value around 0.5. This is a useful relatedness metric when severe loss-of-heterozygosity removes many heterozygous calls from the tumor sample making the traditional relatedness calculation inaccurate.

If a pedigree file is given, Wright’s method of path coefficients^10^ is used to calculate the expected relatedness. These values can then be compared to the relatedness observed from the genotypes. For somatic samples, the user can also specify a “groups” file where sample identifiers appearing on the same line are expected to be identical; for example, three biopsies from each of two individuals would appear as three comma-separated sample identifiers on two separate lines.

Finally, the output is reported both as text and as an interactive HTML page. When using the webpage, the user can toggle which relatedness metrics (IBS0, IBS2, relatedness, homozygous concordance, shared heterozygotes, shared homozygous alternates) to plot for the X and Y coordinates and, if expected groups were given (e.g. tumor-normal pairs) on the command-line, points are colored according to their expected relatedness. This setup means that points of similar colors should cluster together. In addition, *somalier* plots the per-sample output in a separate plot with selectable axes; this functionality allows one to evaluate predicted vs. reported sex and depth across samples.

*Somalier* requires htslib (https://htslib.org). It is written in the Nim programming language (https://nim-lang.org) which compiles to C, and also utilizes our *hts-nim*^11^ library. It is distributed as a static binary and the source code is available at https://github.com/brentp/somalier under an academic license.

## Results

### Glioma patients with 3 samples

We ran *somalier* on BAM alignment files from five individuals, each with three assays: a normal sample, a tumor sample, and cell-free DNA, for a total of 15 samples. The extraction step, which creates the genome sketch and can be parallelized by sample, required roughly three minutes per sample with a single CPU. Once extracted, the *relate* step, which computes the relatedness measures for each sample pair, required less than one second. *Somalier* was able to clearly group samples using the default sites provided with the software (**Figure 2**). Because the site selection is so strict, none of the sample-pairs from the same individual had an IBS0 metric above 0, indicating that those sites are genotyped correctly. The user can specify expected groups of samples (e.g., from the same individual) with sample pairs expected to be identical colored as orange. With this layout that colors sample-pairs by expected relatedness and positions them by observed relatedness (as computed from the genotypes estimated from the alignments), it is simple for the researcher to quickly spot problems. For example, **Figure 2A** illustrates an obvious mix-up where samples expected to be unrelated have a high IBS2 and low IBS0. Since the plot is interactive, the user can then hover over points that appear out of place (in this example, the green points that cluster with the orange) to learn which samples are involved. After correcting the sample manifest based on this observation, and re-running the relatedness calculation, the updated plot shows that all samples cluster as expected given their relatedness. (**Figure 2B**).

**Figure 2.**
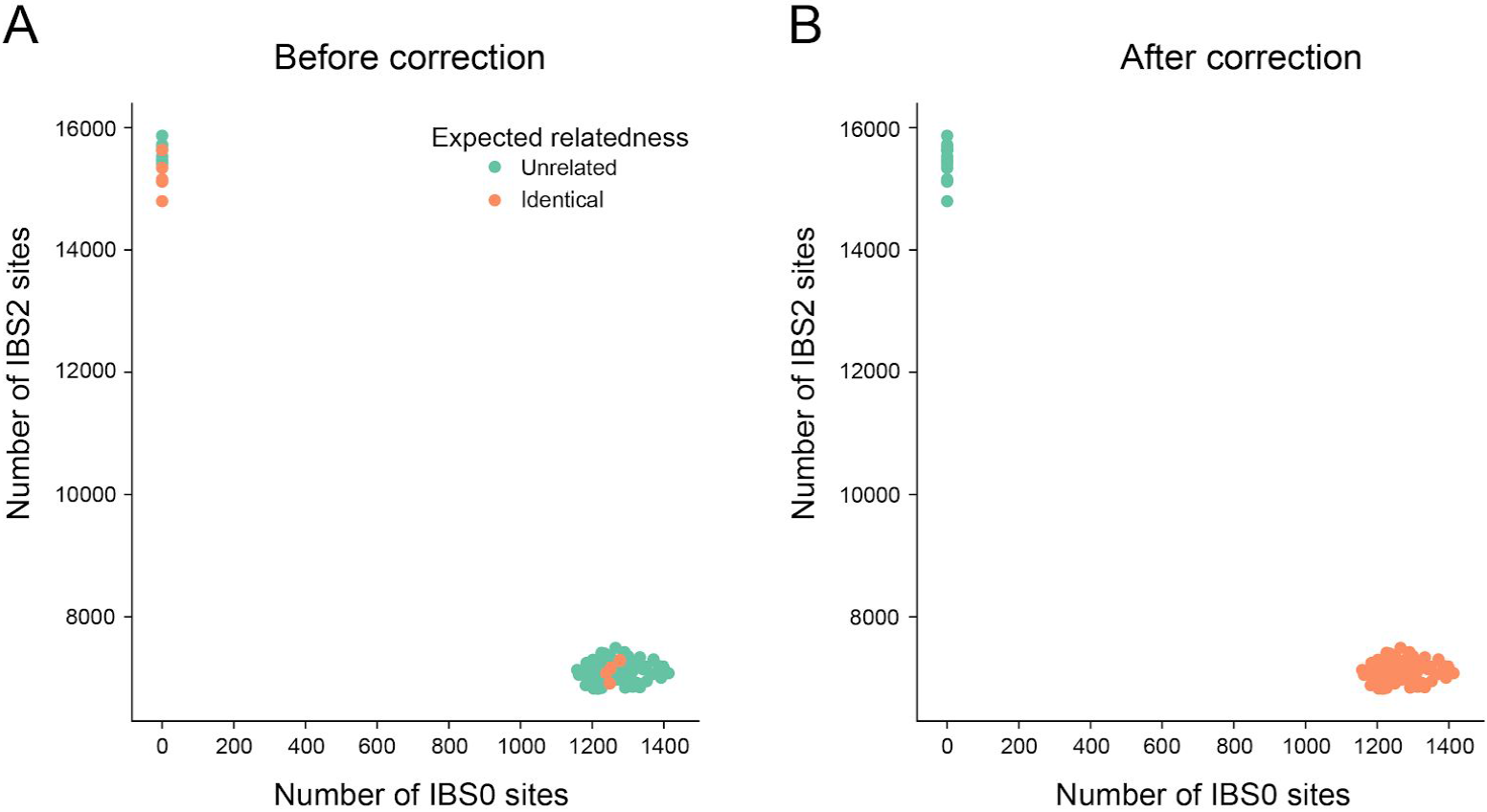
Glioma samples before and after correction. **A.** A comparison of the IBS0 (number of sites where 1 sample is homozygous reference and another is homozygous alternate) and IBS2 (count of sites where samples have the same genotype) metric for 15 samples. Each point is a pair of samples. Points are positioned by the values calculated from the alignment files and colored by whether they are expected to be identical, as indicated from the command-line. In this case, sample swaps are visible as orange points that cluster with green points, and vice versa. The user is able to hover on each point to see the sample-pair involved and to change the X and Y axes to any of the metrics calculated by *somalier*. **B.** An updated version of the plot in panel A after the sample identities have been corrected (per the information provided by panel A) in the manifest after re-running *somalier*.

### 1000 genomes high-coverage samples

In order to evaluate the scalability and accuracy of *somalier*, we used the recently released high-coverage data from 2504 samples in the 1000 Genomes Project (Michael Zody, personal communication). We used the VCF downloaded from: ftp://ftp.1000genomes.ebi.ac.uk/vol1/ftp/data_collections/1000G_2504_high_coverage/working/20190425_NYGC_GATK/ and extracted sites for all 2504 samples. The extraction from VCF completed in about 30 seconds. Comparing each sample against all other samples (a total of 3,133,756 == 2504 * 2503 * 2 comparisons) required merely 9 seconds, following 1.1 seconds to parse the sketches and roughly 2 seconds to write the output. Although the 1000 Genomes Project provides a pedigree file, none of the samples included in the 2504 are indicated to be related by that file. However, using *somalier*, we found 8 apparent parent-child pairs (NA19904-NA19913, NA20320-NA20321, NA20317-NA20318, NA20359-NA20362, NA20334-NA20335, HG03750-HG03754, NA20882-NA20900, NA20881-NA20900) 4 full-sibling pairs (HG02429-HG02479, NA19331-NA19334, HG03733-HG038899, HG03873-HG03998), and 3 second-degree relatives (NA19027-NA19042, NA19625-NA20274, NA21109-NA21135) (**Figure 3**). These same sample pairs also have higher values (as expected) for homozygous concordance. In addition, there are several pairs of samples with a coefficient of relatedness between 0.1 and 0.2 that appear to be more distantly related. An earlier analysis on a different subset of the 1000 Genomes samples uncovered some of these same unreported relationships^12^.

**Figure 3.**
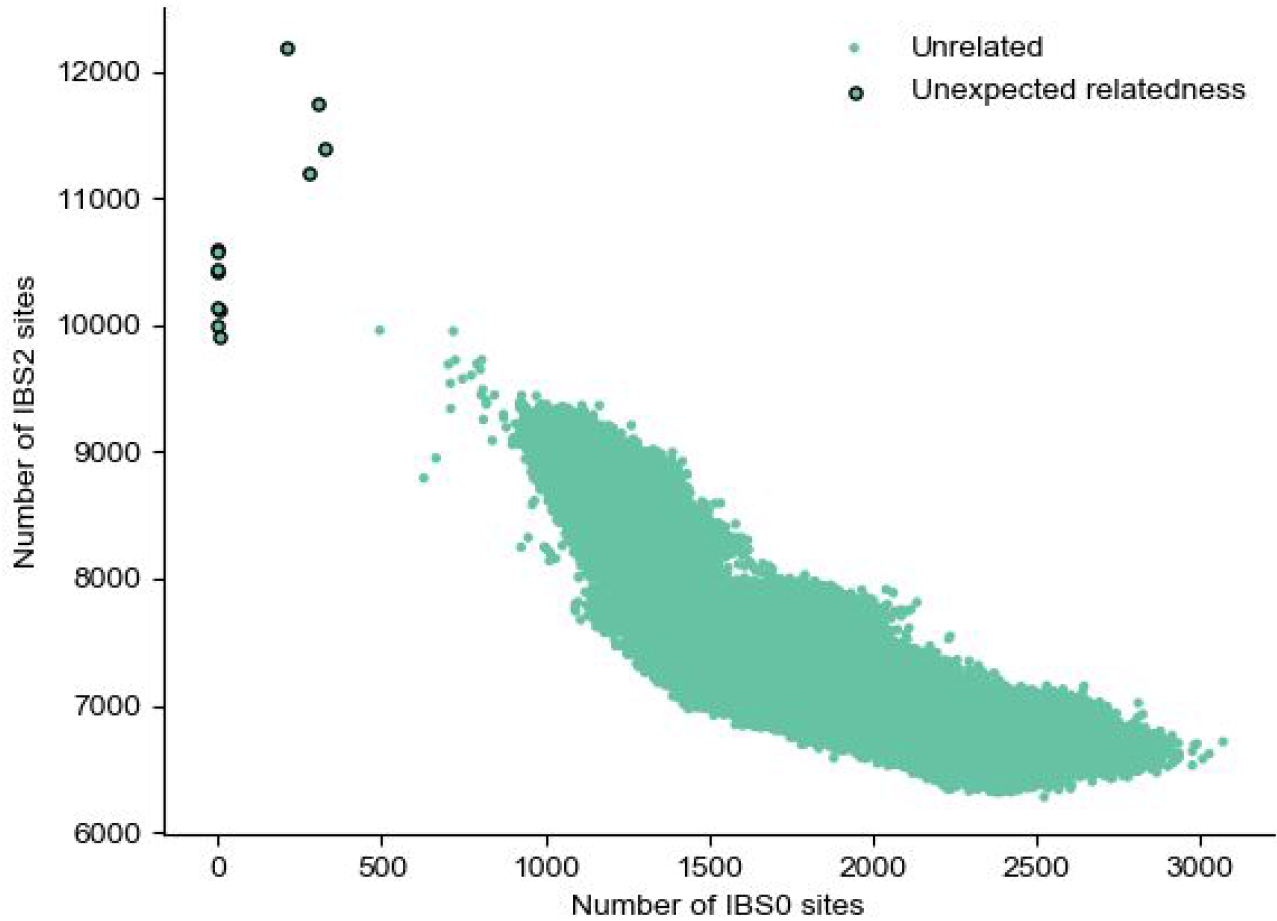
Relatedness plot for thousand genomes samples. Each dot represents a pair of samples. IBS0 on the x-axis is the number of sites where 1 sample is homozygous reference and the other is homozygous alternate. IBS2, on the y axis, is the count of sites where a pair of samples were both homozygous or both heterozygous. Points with IBS0 of 0 are parent-child pairs. The 4 points with IBS0 > 0 and IBS0 < 450 are siblings. There are also several more distantly related sample-pairs.

We also note that several samples indicated to be female in the manifest appear to have lost an X chromosome as they have lower depth and no heterozygous sites (**Figure 4A**). However, they also lack coverage on the Y chromosome (**Figure 4**); as such, we think that loss of X in these cell-line samples is more likely than a sample swap or manifest error.

**Figure 4.**
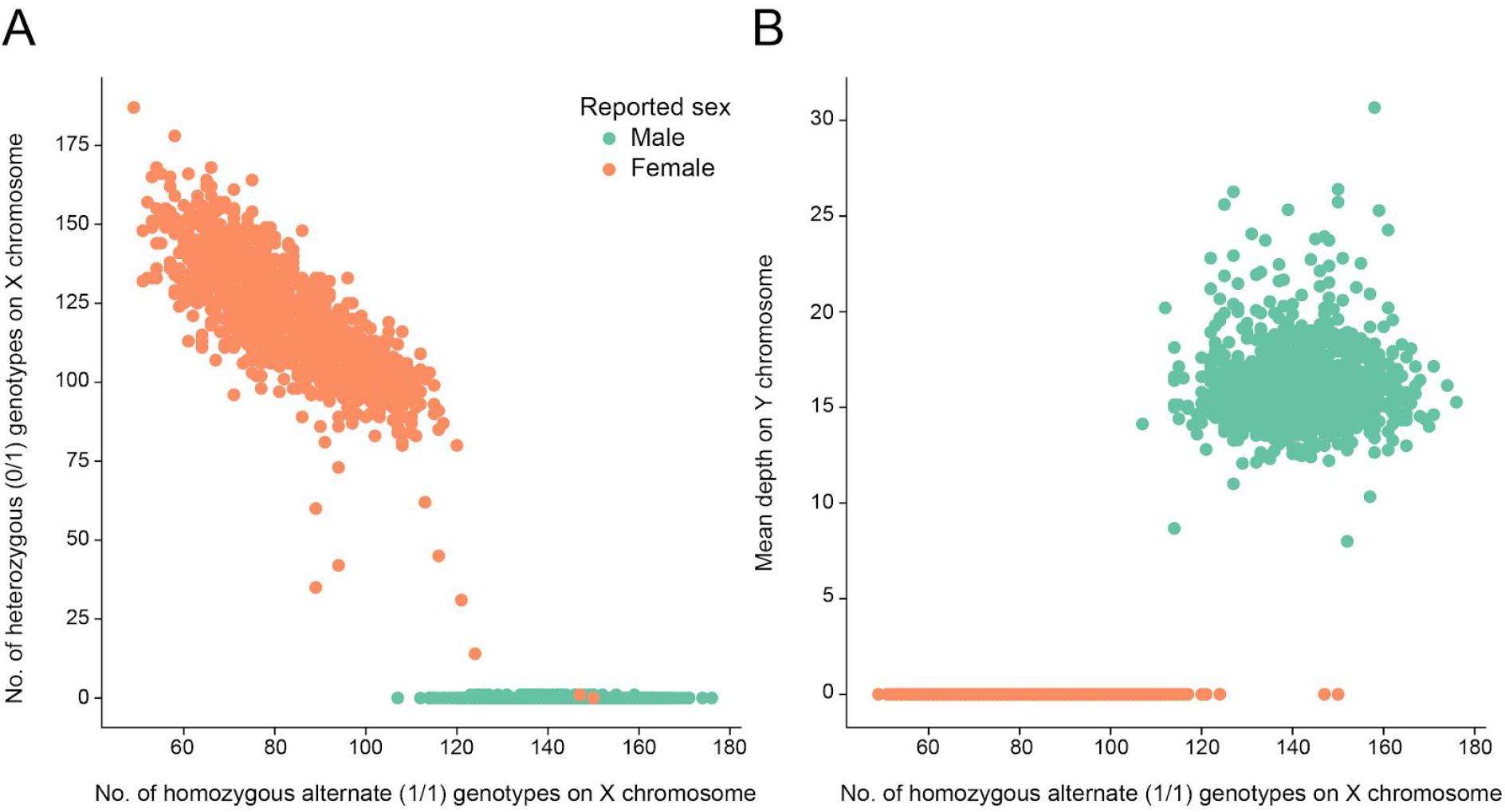
Sex quality control on thousand genomes samples. Each point is a sample colored as orange if the sample is indicated as female and green if it is indicated as male; all data is for the X chromosome. **A**. The number of homozygous alternate sites on the x-axis and the number of heterozygous sites on the y-axis. Males and females separate with few exceptions. **B.** The number of homozygous alternate sites on the x-axis compared to the mean depth on the Y chromosome. Males and females reported in the manifest separate perfectly, indicating that some females may have experienced a complete loss of the X chromosome.

Finally, *somalier* also provides other sample metrics including mean depth, counts of each genotype, mean allele balance, and others that are useful for sample quality control. The user can customize the visualization on the interactive web page by choosing which metrics to display on the X and Y axes.

## Discussion

We have introduced *somalier* to efficiently detect sample swaps and mismatched samples in both cancer and other sequencing projects. On a set of 15 samples, we were able to detect and correct sample swaps using the text and HTML output from *somalier*, which ran in less than a second. In addition, *somalier* can be used to provide an accurate relatedness estimate using *homozygous concordance* even under severe loss-of-heterozygosity. We have designed it to measure relatedness very quickly despite assaying the alignments directly, and we have shown that using a carefully selected set of sites facilitates accurate separation of related from unrelated samples even on a small gene panel.

We have carefully selected the sites assayed by *somalier* to minimize sequencing artefacts and variant calling errors. In addition, we distribute a set of sites for genome build GRCh37 that is compatible with genome build GRCh38. Because the sets are identical, we can compare samples aligned to either genome build. This becomes more important as research groups switch to GRCh38. In fact, in comparing the recently released high coverage 1000 Genomes samples (aligned to GRCh38) to the Simons Diversity Project samples^13^ (aligned to GRCh37), we found several samples shared between these projects. To our knowledge, this has not been previously reported. These findings highlight the utility and novelty of *somalier*, as it enables comparing across large cohorts.

Previous tools such as peddy provide similar functionality when a jointly-called, germline VCF is provided. However, that is often not practical for cancer studies. In addition, HYSYS can detect sample-swaps in cancer samples using homozygous concordance, however it requires a custom text format which reports germline variants that have already been called across all samples. The sketch format used by *somalier* is a simple binary format and we include an example in the repository that demonstrates reading the data in a simple python script and performing ancestry estimation using principal components analysis. While *somalier* can also utilize any number of VCF files as input, we expect that the simplicity and speed of using alignment files will make that the most frequent mode of use.

## Acknowledgements

Thanks to Wayne Clark for noting that the anomaly in the 1000 Genomes samples was loss of chromosome X rather than a sex swap. Thanks to several early users of *somalier* for valuable feedback. The development of somalier was supported by an NIH award to Base2 Genomics, LLC (R41HG010126) and a U24 award to ARQ and GTM.

## Supplement

Sites file for hg38: https://github.com/brentp/somalier/files/3412456/sites.hg38.vcf.gz Sites file for GRCh37: https://github.com/brentp/somalier/files/3412455/sites.GRCh37.vcf.gz

## Notes

#### Summary of Updates

Add Joe to Author list.

https://github.com/brentp/somalier

## References

1. Cibulskis, K. et al. Sensitive detection of somatic point mutations in impure and heterogeneous cancer samples. Nat. Biotechnol. 31, 213–219 (2013).

2. McKenna, A. et al. The Genome Analysis Toolkit: a MapReduce framework for analyzing next-generation DNA sequencing data. Genome Res. 20, 1297–1303 (2010).

3. Pedersen, B. S. & Quinlan, A. R. Who’s Who? Detecting and Resolving Sample Anomalies in Human DNA Sequencing Studies with Peddy. Am. J. Hum. Genet. (2017). doi:10.1016/j.ajhg.2017.01.017

4. Manichaikul, A. et al. Robust relationship inference in genome-wide association studies. Bioinformatics 26, 2867–2873 (2010).

5. Danecek, P. et al. The variant call format and VCFtools. Bioinformatics 27, 2156–2158 (2011).

6. Bergmann, E. A., Chen, B.-J., Arora, K., Vacic, V. & Zody, M. C. Conpair: concordance and contamination estimator for matched tumor-normal pairs. Bioinformatics 32, 3196–3198 (2016).

7. Lek, M. et al. Analysis of protein-coding genetic variation in 60,706 humans. Nature 536, 285–291 (2016).

8. Li, H. Toward better understanding of artifacts in variant calling from high-coverage samples. Bioinformatics 30, 2843–2851 (2014).

9. Schröder, J., Corbin, V. & Papenfuss, A. T. HYSYS: have you swapped your samples? Bioinformatics 33, 596–598 (2017).

10. The Method of Path Coefficients on JSTOR. Available at: https://www.jstor.org/stable/2957502. (Accessed: 10th October 2019)

11. Pedersen, B. S. & Quinlan, A. R. hts-nim: scripting high-performance genomic analyses. Bioinformatics 34, 3387–3389 (2018).

12. Roslin, N. M., Weili, L., Paterson, A. D. & Strug, L. J. Quality control analysis of the 1000 Genomes Project Omni2.5 genotypes. bioRxiv 078600 (2016). doi:10.1101/078600

13. Mallick, S. et al. The Simons Genome Diversity Project: 300 genomes from 142 diverse populations. Nature 538, 201–206 (2016).

